# Crosstalk between Rac and Rho GTPase activity mediated by Arhgef11 and Arhgef12 coordinates cell protrusion-retraction cycles

**DOI:** 10.1101/2023.02.20.529203

**Authors:** Suchet Nanda, Abram Calderon, Thanh-Thuy Duong, Johannes Koch, Arya Sachan, Xiaoyi Xin, Djamschid Solouk, Yao-Wen Wu, Perihan Nalbant, Leif Dehmelt

## Abstract

Rho GTPase crosstalk is thought to play a key role in the spatio-temporal coordination of cytoskeletal dynamics during cell migration. Here, we directly investigated crosstalk between the major Rho GTPases Rho, Rac and Cdc42 by combining acute activity perturbation with activity measurements in individual, mammalian cells. As expected for their proposed mutual inhibition, we confirmed that Rho inhibits Rac activity. However, surprisingly, we found that Rac strongly stimulates Rho activity. We hypothesized that this crosstalk might play a role in mediating the tight spatio-temporal coupling between cell protrusions and retractions that are typically observed during mesenchymal cell migration. Using new, improved activity sensors for endogenous Rho GTPases, we find that Rac activation is tightly and precisely coupled to local cell protrusions, followed by Rho activation during retraction. In a screen for potential crosstalk mediators, we find that a subset of the Rho activating Lbc-type GEFs, in particular Arhgef11 and Arhgef12, are enriched at transient cell protrusions and retractions. Furthermore, via an optogenetic approach, we show that these Lbc GEFs are recruited to the plasma membrane by active Rac, suggesting that they might link cell protrusion and retraction by mediating Rac/Rho activity crosstalk. Indeed, depletion of these GEFs impaired cell protrusion-retraction dynamics, which was accompanied by a decrease in migration distance and an increase in migration directionality. Thus, our study shows that Arhgef11 and Arhgef12 facilitate effective exploratory cell migration by coordinating the central cell morphogenic processes of cell protrusion and retraction by coupling the activity of the associated small GTPases Rac and Rho.

## Introduction

Cytoskeletal dynamics drive force-generating mechanisms that control changes in cell shape during cell morphogenesis and cell migration (Fletcher & Mullins, 2010). These force-generating mechanisms, in turn, are controlled by signaling networks that are regulated in space and time (Hodge & Ridley, 2016). Members of the Rho GTPase family play important roles in this process. Protrusive forces at the leading edge of cells are typically induced by the Rho GTPase family members Rac1 and Cdc42, which promote nucleation and polymerization of actin filaments and associated proteins (Ridley, 2015). Rac1 induces flat cell protrusions, called lamellipodia, while Cdc42 induces pointed protrusions, called filopodia. RhoA, a related Rho GTPase family member, induces stress fibers that generate contractile forces that originate from Myosin 2 motor mini-filaments that act on anti-parallel actin filaments (Ridley, 2015).

Crosstalk between Rho GTPases appears to play an important role in the coordination of their activities in space and time (Guilluy et al., 2011). In a simple concept, mutually inhibitory crosstalk between Rac1/Cdc42 and RhoA was proposed to spatially segregate cell protrusion and cell contraction between the protrusive, leading edge and the contractile, trailing rear, respectively (Guilluy et al., 2011). Studies using Rho activity sensors in small migrating cells, such as neutrophils, have produced results that are consistent with this idea (Wong et al., 2006; Yang et al., 2016). However, in larger migrating cells, such as fibroblasts, protrusive Rac1/Cdc42 and contractile RhoA signals were both detected near the leading edge of migrating cells, suggesting a more complex relationship between these Rho GTPases (Kraynov et al., 2000; Machacek et al., 2009; Nalbant et al., 2004; Pertz et al., 2006). Furthermore, mesenchymal cells typically generate highly dynamic regular or irregular cycles of cell protrusion and retraction near the leading cell edge, which are thought to play an exploratory role in cell migration (Nalbant & Dehmelt, 2018). Thus, in such cells protrusion and retraction dynamics appear to be tightly coupled in space and time. Therefore, specific mechanisms must exist that generate these highly dynamic cycles of cell protrusion and retraction.

Recent studies revealed transient activity pulses or traveling waves of signal network components that promote cell protrusion, including Cdc42, actin nucleation promoting factors (NPFs), and Ras-type GTPases (Huang et al., 2013; Miao et al., 2017; Weiner et al., 2007; Yang et al., 2016). These activity patterns are typical for signal networks that combine both positive and negative feedback regulation to generate excitable or oscillatory system dynamics (Tyson et al., 2003). We and others recently found that the contractility-inducing GTPase Rho is a central component of a signal network that generates pulses and waves of cell contraction (Bement et al., 2015; Graessl et al., 2017; Kamps et al., 2020). Here we hypothesize, that a link between these excitable or oscillatory signal networks might play a role in coordinating the highly dynamic cycles and bursts of cell protrusion and cell retraction that are observed in exploratory, mesenchymal cell migration.

Previous observations of Rho GTPase crosstalk have relied on slow perturbations and/or indirect evidence. Here, we combine acute activity perturbation techniques based on chemically-induced dimerization (CID) as well as optogenetic approaches with continuous monitoring of response dynamics to directly investigate Rho GTPase crosstalk in living cells. Importantly, the perturbation constructs that were used in our studies are based on constitutively active Rho GTPase mutants (Liu et al., 2014; Wu et al., 2009), which are not prone to potential feedback regulation and adaptive responses. We thereby minimize secondary effects, which might otherwise mask cause and effect relationships. As expected from their proposed mutual inhibition, acute activation of RhoA led to a decrease in Rac activity. Surprisingly, our studies revealed robust activation of Rho after acute Rac1 activation, which is contrary to the expected mutual inhibition between these signals. Furthermore, by using novel, improved sensors to measure endogenous Rac1 and Rho activity, we reveal a tight correlation between increased, local Rac activity at the plasma membrane during cell protrusion and increased Rho activity during cell retraction.

Investigations into the molecular mechanisms revealed that the Rho-activating Lbc-type GEFs Arhgef11 and Arhgef12 act as Rac effectors and thereby can mediate the observed Rac/Rho activity crosstalk. Furthermore, our data shows that Arhgef11 and Arhgef12 are required for effective spatio-temporal coupling between cell protrusion and retraction dynamics, and that this is critical for efficient exploratory cell migration.

## Results

### Analysis of Rho GTPase crosstalk in living cells

To directly investigate Rho GTPase crosstalk, we sought to develop methods that enable the combination of acute activity perturbations and simultaneous activity monitoring in individual, living cells. To induce such acute perturbations, we extended an approach that we developed previously (Liu et al., 2014), which is based on reversible, chemically-induced plasma membrane targeting of constitutively active Rho GTPase mutants (Figure 1A). Analogous to the published approach (Liu et al., 2014), we removed the C-terminal CAAX motif found in Rho GTPases, which normally acts as a membrane anchor after post-translational geranylgeranylation (Katayama et al., 1991; Ménard et al., 1992; Ziman et al., 1993). Instead, we added the FKBP’ heterodimerization domain. This allowed us to induce robust, reversible targeting of Rho GTPases from the cytosol to the plasma membrane, where they interact with a plasma membrane anchored heterodimerization partner (eDHFR) upon addition of a chemical dimerizer (SLF’-TMP). In our previous study, we found that the recruitment of constitutively active Rac1 induced robust and reversible formation of lamellipodia, which are a typical phenotype of increased Rac1 activity (Liu et al., 2014). Here, we extended this perturbation strategy to the three best-characterized Rho GTPases Rac1, Cdc42, and RhoA. As Neuro-2a neuroblastoma cells show robust morphological changes in response to active Rac1 (Schonichen et al., 2013), we used this cell line for our initial analyses. As expected, Rac1 and Cdc42 plasma membrane targeting induced the formation of cell protrusions, while RhoA plasma membrane targeting induced cell contraction (Figure 1B and Movie 1). Inhibition of plasma membrane targeting with a small molecule competitor (TMP) reversed the Rho GTPase-induced phenotypes (Figure 1B and Movie 1).

**Figure 1:**
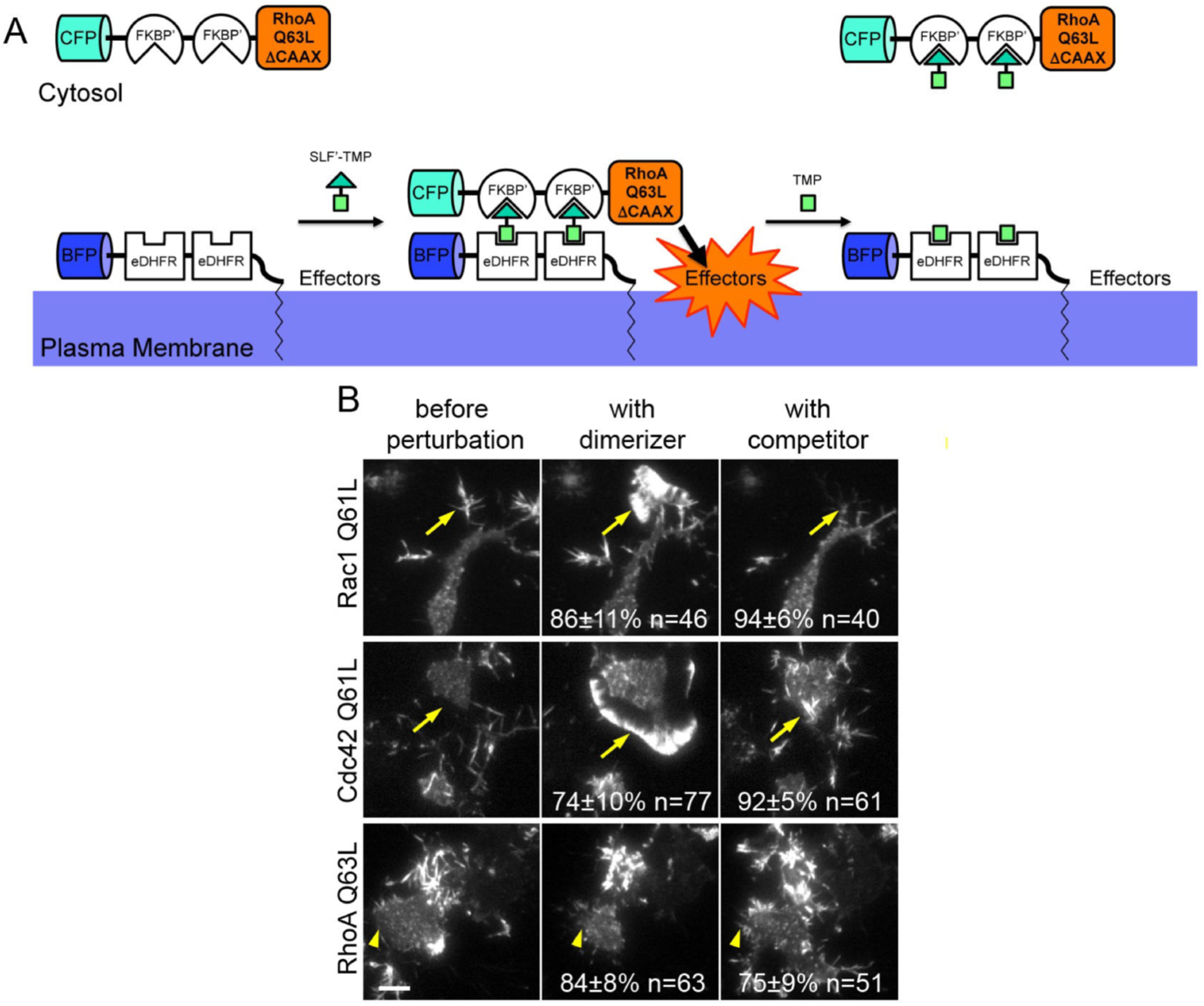
A general method for acute perturbation of Rho GTPase activity in living cells. **A:** Schematic of acute, reversible Rho GTPase activity perturbation via chemically-induced dimerization. **B:** Representative frames from TIRF microscopy time series of mCherry-Actin obtained 30s before, 24 min during and 24min after acute Rho GTPase activation in Neuro-2a neuroblastoma cells (see also Movie 1). Yellow arrows point to cell areas that reversibly generate lamellipodial protrusions during Rac1 or Cdc42 activation, and yellow arrowheads point to areas that undergo reversible retraction during RhoA activation. Observations are representative for 3 independent repetitions with a total of at least 40 cells per condition (exact numbers of cells are indicated in individual panels). Numbers in middle panels indicate percentage ± standard error of the mean of cells that initiate protrusion (Rac1/Cdc42) or retraction (RhoA) after addition of dimerizer. Numbers in right panels indicate percentage of reacting cells that showed a phenotypic reversal. Scale bar: 10 μm. CFP: cyan fluorescent protein; BFP: blue fluorescent protein; FKBP’: FK506-Binding Protein with F36V mutation; eDHFR: *E. coli* dihydrofolate reductase; SLF’: synthetic ligand of FKBP’; TMP: eDHFR interacting small molecule trimethoprim.

Next, we combined acute Rho GTPase perturbations with Rho GTPase activity measurements. To measure Rho, Cdc42, and Rac activity, we used TIRF microscopy-based sensor constructs that we developed previously (Graessl et al., 2017). These Rho GTPase activity measurements are based on the plasma membrane recruitment of GTPase binding domains (GBDs) from specific effector proteins: Rhotekin, WASP, and p67^phox^, respectively. While these effector domains are known to be specific for the respective Rho, Cdc42, and Rac subgroups, they are not expected to distinguish between more closely related family members. The Rhotekin-based Rho sensor will detect active RhoA, RhoB, and RhoC, the WASP-based Cdc42 sensor will detect active Cdc42, TC10, and TCL and the p67^phox^-based Rac sensor will detect Rac1, Rac2, and Rac3. The effectors and cellular functions of these closely related GTPases are very similar. Therefore, we consider their combined activity for our crosstalk analyses and refer to sensor measurements using the corresponding Rho GTPase subfamily names Rac, Cdc42, and Rho. During the time course of dimerizer induced activity perturbations, significant changes in cell volume can occur, which could also change the TIRF signal. To control for this potential artifact, we co-transfected a cytosolic cell filler that acts as a control construct and used this to correct changes in the sensor signal (see Methods). Figure 2B shows the Rac1 perturbation and the raw, uncorrected Rho activity sensor and control sensor response measurements in a representative cell (see also Movie 2). Here, an increase of the Rho sensor response was accompanied by a decrease in the control sensor signal, showing that the uncorrected sensor measurement underestimated the actual response.

**Figure 2:**
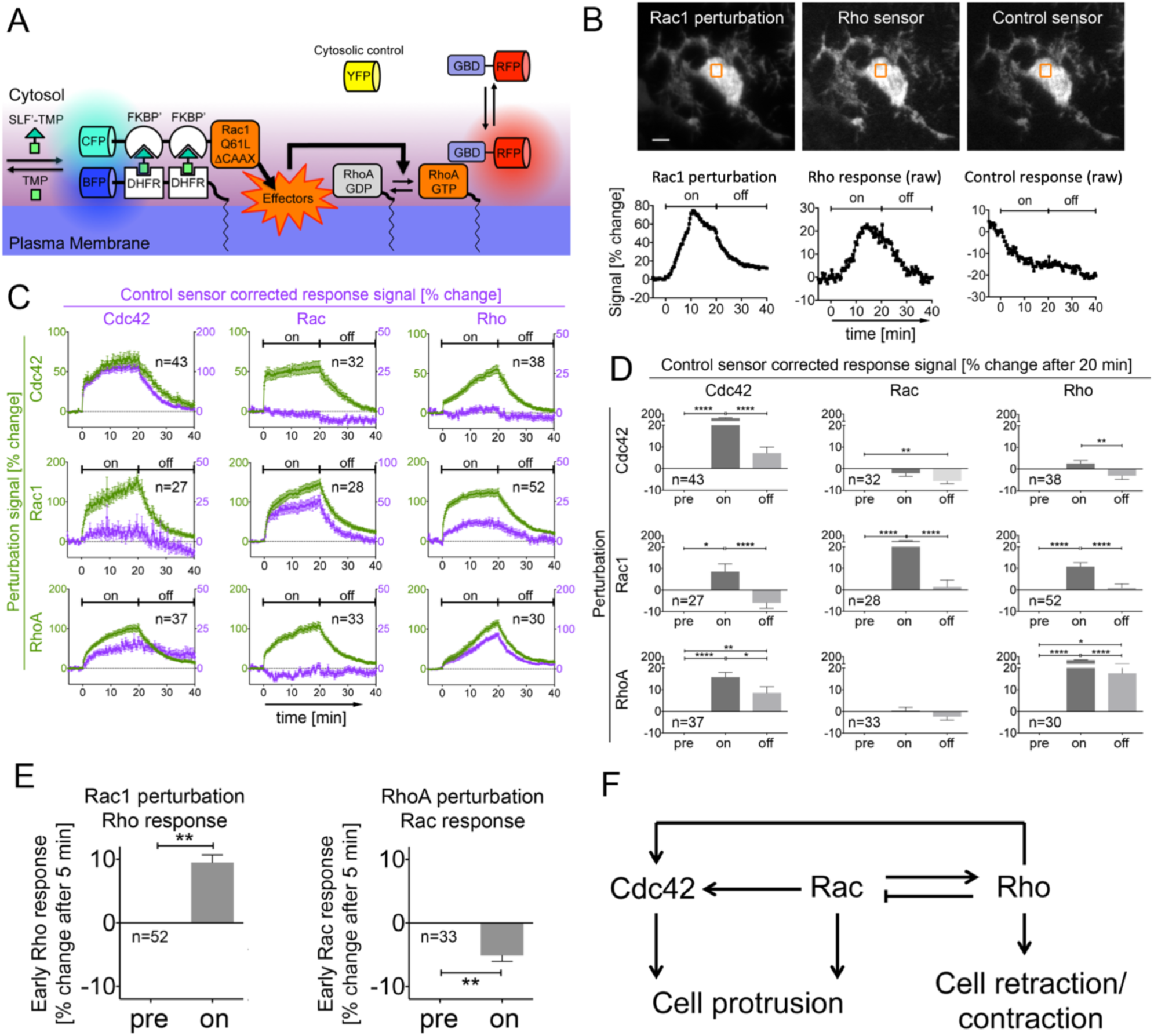
Analysis of global Rho GTPase crosstalk in living cells. **A:** Schematic of acute activity perturbation and combined activity measurement strategy. **B-E:** Analysis of perturbation-response relationships of Rho GTPase activity in Neuro-2a cells. **B:** Representative TIRF images before dimerizer addition (top) and Rac1 perturbation and uncorrected, raw Rho sensor and raw control sensor signal kinetics (bottom) corresponding to orange boxes (see also Movie 2). All constructs are predominantly cytosolic and homogenously distributed in the cell bodies and neurite-like protrusions. **C:** Average perturbation and control-corrected activity sensor signal kinetics for all crosstalk combinations. **D:** Statistical analysis of average sensor signal changes during and after Rho GTPase activity perturbation. For these analyses, responses at time points before (pre), and 20 minutes after addition of dimerizer (on) or competitor (off) were considered. **E:** Statistical analysis of average sensor signal changes during acute Rho GTPase activity perturbation. Responses at time points before (pre), and 5 minutes after dimerizer addition (on) are shown. **F:** Influence diagram that summarizes significant activity response measurements at 5 minutes after dimerizer addition. All observations and measurements are based on at least 3 independent repetitions with a total of at least 27 cells per condition (exact numbers of cells are indicated in individual panels). Scale bars: 5 µm. *: P<0.05; **: P<0.01; ****: P<0.0001; D: One-way ANOVA; E: Student’s t-Test. Error bars represent standard error of the mean.

Several crosstalk measurements were in agreement with previous observations. As expected, the strongest response in the activity for a particular sensor was observed upon activation of the corresponding Rho GTPase (Figure 2C-D). In a previous study (Graessl et al., 2017), we observed that Rho activity pulses precede increased Cdc42 activity with a delay of 10.1±1.8 s, suggesting that Cdc42 might be activated downstream of Rho activity. However, these previous observations only indicated a correlation between Rho and Cdc42 and did not show a causal relationship between these activities. Here we found that acute activation of RhoA indeed stimulates Cdc42 activity (Figure 2C-D). We also observed inhibition of Rac activity by acute

RhoA activation, which is in agreement with the previously proposed mutual inhibition between these GTPases (Figure 2C,E) (Guilluy et al., 2011; Kuo et al., 2011; Ohta et al., 2006; Sanz-Moreno et al., 2008; Vicente-Manzanares et al., 2011). Interestingly, in our studies, activation of Rho was the strongest and most robust crosstalk after activation of Rac (Figure 2B-E and Movie 2). This was surprising as it contradicts the proposed mutual inhibitory relationship between RhoA and Rac1 (Guilluy et al., 2011). Finally, we observed significant activation of Cdc42 by Rac1, which could result either from direct crosstalk between these GTPases, or from indirect activation via Rho (Figure 2C-D, Figure 2F)

As the observation that Rac1 robustly activates Rho was unexpected, we sought to further investigate this relationship with an alternative method. Therefore, we used the light-controlled PA-Rac1 construct which was previously developed by fusing a LOV2 domain to constitutively active Rac1 (Wu et al., 2009). After illumination with a wavelength of 445nm, this LOV-domain quickly unfolds within milliseconds and unblocks active Rac1, which can then interact with its effectors (Wu et al., 2009). In the absence of 445nm light, the LOV2 domain refolds within several seconds, which again blocks the associated active Rac1 (Graessl et al., 2017; Wang et al., 2016; Wu et al., 2009). By combining this method with Rho activity measurements (Figure 3A), we confirmed Rac1-dependent Rho activation in Neuro-2a cells (Figure 3B). In contrast, very little change in signal was observed using the control sensor (Figure 3B).

**Figure 3:**
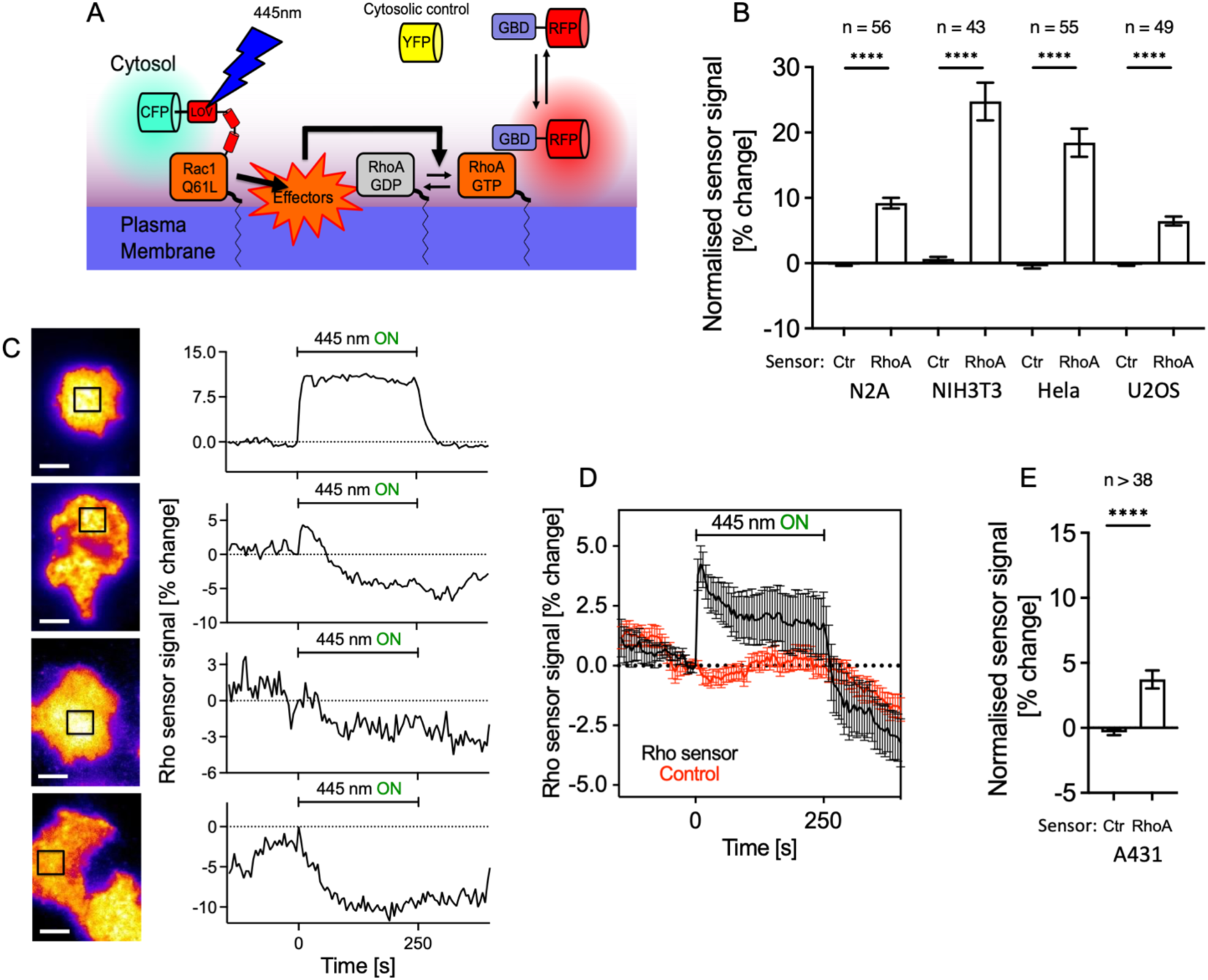
Rac/Rho crosstalk is commonly observed in adherent mammalian cell lines and can trigger a dynamic Rho activity response. **A:** Scheme of optogenetic approach to measure Rac/Rho activity crosstalk. **B-E**: Measurement of Rho activity dynamics during acute, optogenetic activation of Rac. **B, E**: Rho activity response to acute Rac1 perturbation in commonly used cell lines. The difference between 5 measurements before and 5 measurements after the onset of illumination is shown, each corresponding to a time frame of 25s. **C**: Typical Rho activity response dynamics observed in A431 cells. Left: representative TIRF images from video microscopy time series. Right: Rho activity dynamics corresponding to black boxes in left panels. 67% of cells showed a discernible, reversible Rho activity response. Of these responding cells, 29% showed a continuous Rho activation during Rac1 photoactivation (top panels), 38% of the cells showed a single, transient activity pulse response and 25% showed no response (middle panels). 8% of cells showed a negative response (bottom panels). **D**: Measurement of average Rho activity sensor kinetics before, during and after Rac activation in A431 cells. ****: P<0.0001; Student’s t-Test. Error bars represent standard error of the mean. Scale bars: 10µm.

We next sought to determine whether Rac1-induced Rho activation was unique to Neuro-2a neuroblastoma cells. Therefore, we repeated the photoactivation experiment in a panel of commonly used cell lines, including NIH-3T3, HeLa, U2OS and A431 cells. Similar as in Neuro-2a cells, we observed a robust activation of Rho during acute, continuous, Rac1 activation in these cells (Figure 3B). However, while Neuro-2a, NIH-3T3 and HeLa cells predominantly responded with a constant, increased Rho activity level, U2OS (Supplementary Figure 1) and A431 cells (Figure 3C-E), responded more dynamically. This more dynamic response suggests additional regulatory mechanisms, for example via positive or negative feedback loops.

### Rac and Rho activity dynamics in cell protrusion-retraction cycles

The observed stimulation of transient Rho activity dynamics by Rac1 in U2OS and A431 cells might play a role in cellular processes that are characterized by transient cell shape changes. We were particularly intrigued by the idea that the Rac1/Rho crosstalk might play a role in coupling these activities during cell protrusion-retraction cycles. A431 cells are well known to generate transient cell protrusions that are followed by transient cell retractions both after growth factor stimulation (Gagliardi et al., 2015) and spontaneously (Clark et al., 2022). To study a potential role of Rho GTPase crosstalk in this system, we first characterized the spatio-temporal coupling of Rac and Rho activity with cell shape changes. Initial experiments using the sensors described above showed only weak signals. This suggests that these sensors are not sensitive enough to detect the activation of Rho GTPases during spontaneous cell morphodynamics. To increase their sensitivity, we generated constructs that contain tandem GTPase binding domains that could benefit from binding multiple GTPase molecules at the same time. This alternative sensor design might lead to increased interactions with active GTPases due to increased avidity. As these constructs can compete with endogenous effectors, we expressed them at very low levels, using the truncated delCMV promotor and detected their localization using highly sensitive TIRF microscopy equipment.

Interestingly, we observed very tight spatio-temporal coupling between Rac activity and cell protrusion and Rho activity and cell retraction (Figure 4A, E, see also Movies 3, 4 and 5). To quantify the coupling between Rho GTPase activity and cell shape changes, we used the open-source ADAPT ImageJ plugin (Barry et al., 2015), which is well-suited to track cell edge movements and associated fluorescence signals at the cell edge (Figure 4B). We first modified the plugin to optimize extraction of our sensor signals. In the original implementation, regions are investigated that extend equally both inside and outside the cell border. We changed these analysis areas to only include signals approximately 3 μm towards the inside of the cell border to increase the signal measurement sensitivity (see methods for details). We then used this modified plugin to extract spatio-temporal fluorescence signal and cell edge velocity maps (Figure 4C) and temporal signal-cell edge velocity cross-correlation functions (Figure 4F). The measured correlation between Rac and cell edge velocity supports the observation that Rac activity increases during cell protrusion (see also Movies 3 and 4). Conversely, the anti-correlation measured for Rho supports the observation that Rho activity increases during cell retraction (negative cell edge velocity, see also Movie 5). Furthermore, the maxima and minima of these functions are only minimally shifted in time, showing that the correlation between sensor signal and cell shape changes does not have a significant delay.

**Figure 4:**
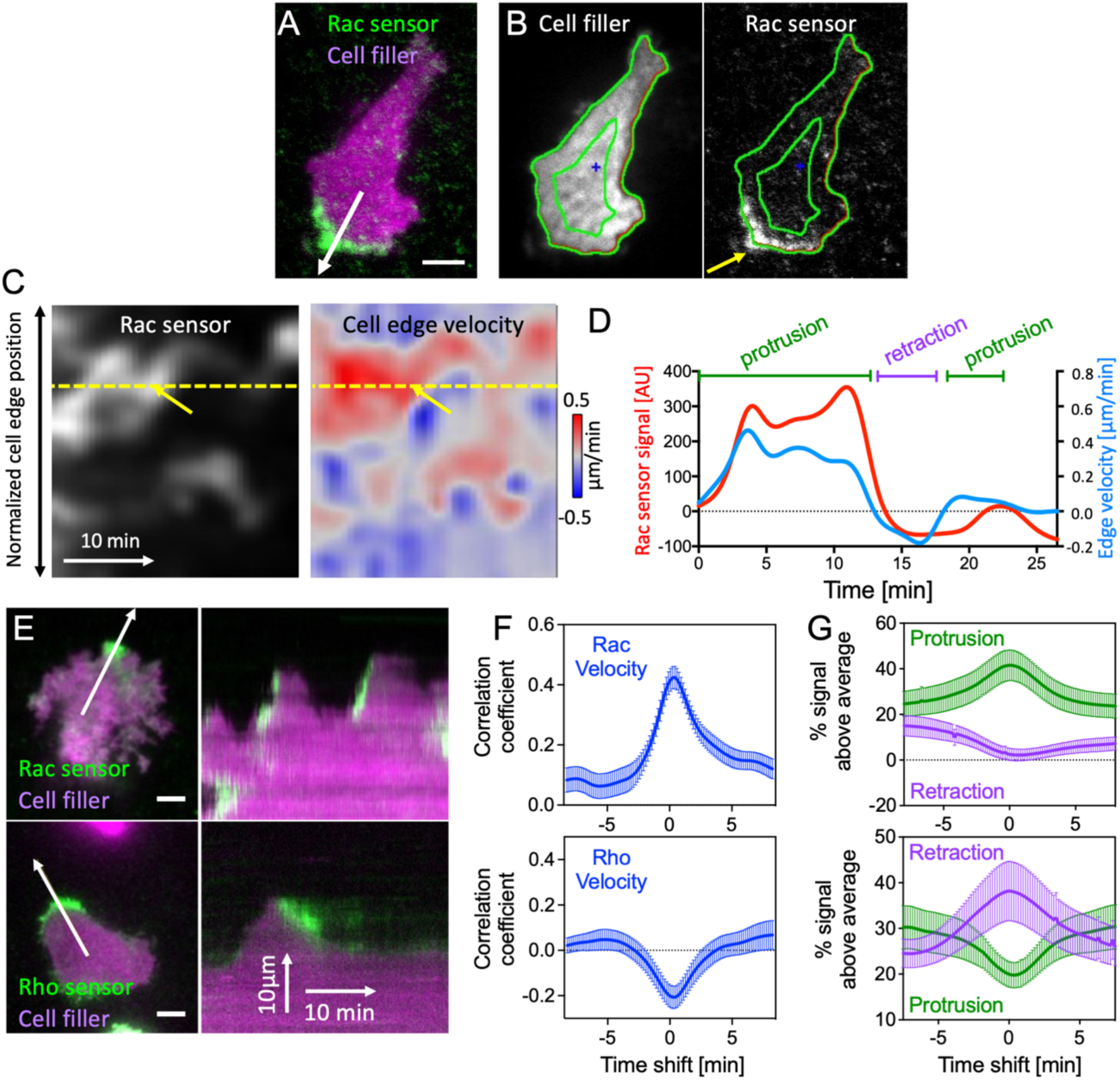
Sequential Rac and Rho activation is tightly coupled to cell protrusion-retraction cycles in space and time. **A**: Representative A431 cell expressing a cytosolic cell filler (mCitrine) and an improved Rac activity sensor (mCherry-3xp67^phox^GBD; top; see also Movie 3). The white arrow marks the direction of a local cell protrusion. **B**: Automated tracing of the cell border using a modified version of the ADAPT plugin (Barry et al., 2015). **C**: Maps generated by the ADAPT script that represent the spatio-temporal dynamics of the Rac sensor signal between the green lines in **B** (left) and of the cell edge velocity (right). The yellow arrows in **C** point to the local protrusion that occurs at the position of the cell area marked by the yellow arrow in **B**. Red areas in the velocity map correspond to local cell protrusions, blue areas to local cell retractions. **D**: Plot of Rac sensor signals and cell edge velocity corresponding to the yellow dotted line in **C**. **E**: Representative TIRF images (left) of A431 cells that generate spontaneous protrusion-retraction cycles and express the Rac or Rho GTPase activity sensors and the cell filler (see also Movies 4 and 5). White arrows represent the protrusion direction. Kymographs (right) correspond to white arrows in TIRF images. **F**: Crosscorrelation between Rac sensor signal and cell edge velocity plotted against the time shift between these measurements. **G**: Enrichment of Rac and Rho sensor signals in protrusions (>0.075μm/min) and retractions (<-0.075μm/min). Values are normalized to average control sensor enrichment measurements. n=3 independent experiments with >21 cells per condition. Error bars represent standard error of the mean. Scale bars: 10µm.

### Development of a new analysis approach to quantify Rac and Rho activity enrichment at the cell border

The correlation functions contain limited information about the relation between the sensor signals and cell shape changes. First, they cannot be used to distinguish between a signal increase during protrusion and a signal decrease during retraction. Both of these events would result in a positive cross-correlation value. However, cell protrusion and cell retraction are very distinct cellular processes that involve distinct sets of regulators. Therefore, measurements that mix these processes such as the signal/cell edge velocity correlation shown in Figure 4F, blur the information specific to these distinct processes. Second, the correlation functions do not provide a measure for the strength of the signal, and how much it is enriched in a particular area of the cell. To overcome these limitations, we developed our own, extended analysis approach, in which we used the signal and velocity maps (Figure 4C) to measure, how much the fluorescence signal is enriched within local cell protrusions or retractions relative to the average signal of the whole cell (Figure 4D, see methods for details). Analogous to temporal cross-correlation functions, we introduced time shifts between the protrusion-retraction events and signal measurements, to obtain temporal signal enrichment functions that show how much the fluorescence signal is enriched or depleted relative to the time periods of protrusion and retraction (Figure 4G).

Applying this analysis to the cell protrusion regulator Rac1 yielded a very clear representation of the observed dynamics (Figure 4G; top panel), i.e. that Rac1 is highly enriched during cell protrusion and slightly depleted during retraction. Conversely, the cell retraction regulator Rho was highly enriched during cell retraction and depleted during protrusion (Figure 4G; bottom panel, see also Movie 5). The enrichment of the activity signal is calculated in percent relative to the average signal of the whole cell attachment area, which is an easily interpretable measure for local sensor signal strength.

Together, these observations show that the dynamic protrusion-retraction cycles that are observed in A431 cells are tightly coupled to corresponding activations of Rac and Rho. Based on these observations, we hypothesized that the Rac-dependent activation of Rho, which we identified in this study, might play a role in the tight coupling of signals in dynamic protrusion-retraction cycles.

### Molecular mechanism of sequential Rac/Rho activity dynamics

To investigate this hypothesis, we sought to identify potential mediators of Rac-dependent Rho activation. A previous biochemical study suggested that Rho activators of the Lbc GEF family might act as effectors of active Rac (Figure 5A) and thereby might mediate this activity crosstalk (Dada et al., 2017). In this hypothetical mechanism, active Rac1 would recruit an Lbc GEF to the plasma membrane, thereby concentrating the GEF at this site to stimulate local Rho activity. To narrow down the set of potential candidates, we investigated the plasma membrane association dynamics of all Lbc type GEFs in relation to cell protrusion and retraction dynamics. In this focused screen, we found that a subset of the Lbc GEFs: Arhgef11 (PDZ-RhoGEF) and Arhgef12 (LARG), were highly enriched at the edge of the cell during protrusion and retraction (Figure 5B-C, see also Movies 6 and 7).

**Figure 5:**
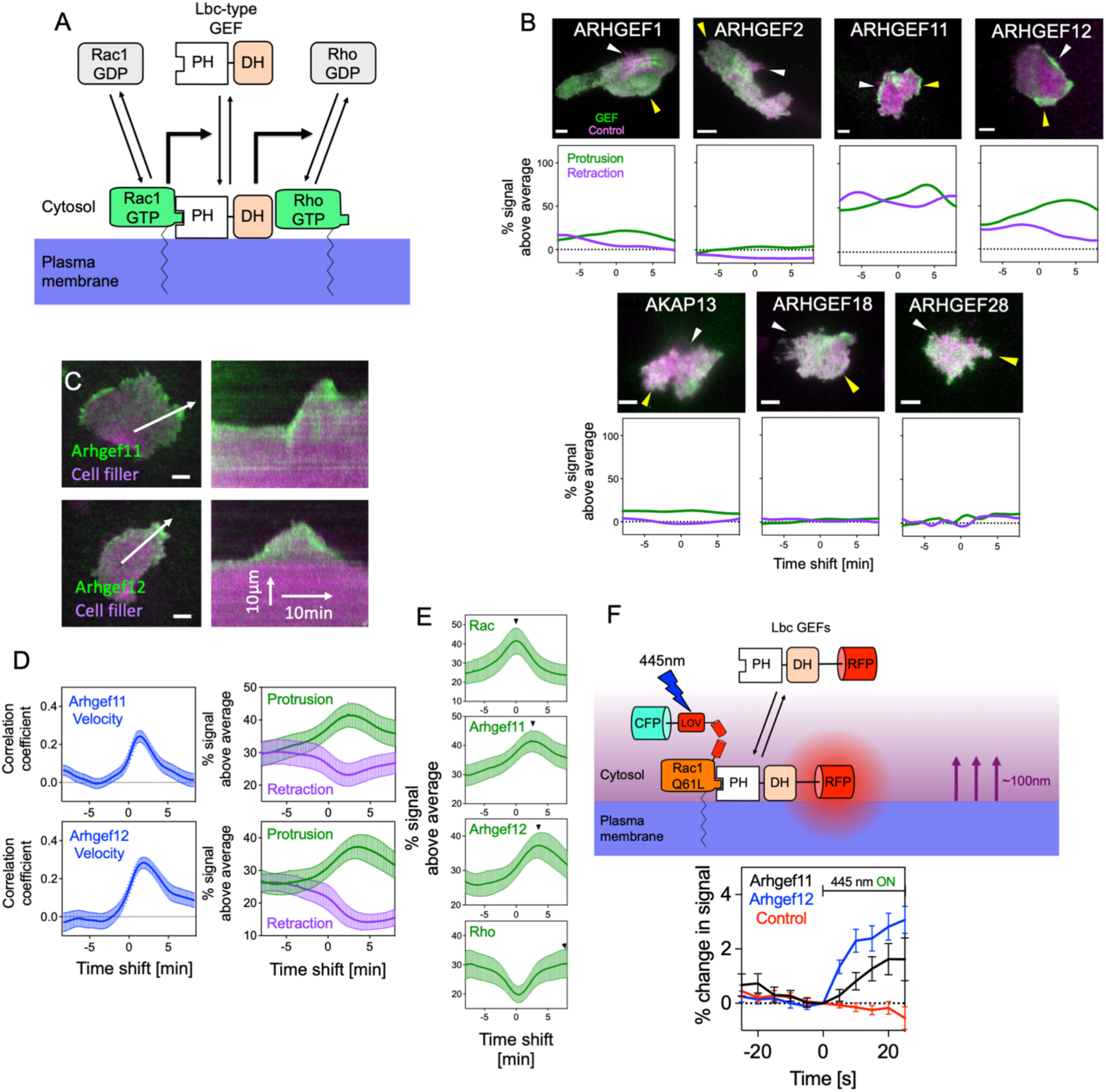
Identification of Arhgef11 and Arhgef12 as Rac effectors in local cell protrusion-retraction cycles. **A:** Schematic representation of a hypothetical mechanism, by which Lbc-type GEFs could mediate Rac1/Rho activity crosstalk. **B**: TIRF microscopy images (top panels) and protrusion-retraction enrichment functions (bottom panels) for representative cells that express Lbc-type GEFs (CMV-GEF, green) and a cytosolic cell filler that acts as a control construct (delCMV-mCitrine, magenta). White and yellow arrows point to local cell retractions and protrusions, respectively. **C**: Representative TIRF images (left) of A431 cells that generate spontaneous protrusion-retraction cycles and express Arhgef11 and Arhgef12 fused to mCherry and the cytosolic cell filler (mCitrine, see also Movies 6 and 7). White arrows represent the protrusion direction. Kymographs (right) correspond to white arrows in TIRF images. **D**: Crosscorrelation (left) between Arhgef11/Arhgef12 signals and cell edge velocity plotted against the time shift between these measurements, and enrichment (right) of Arhgef11/Arhgef12 signals in protrusions and retractions. Arhgef11/Arhgef12 enrichment values are normalized to average control construct enrichment measurements. n=3 independent experiments with >22 cells per condition **E**: Direct comparison of signal enrichment of active Rac, Arhgef11, Arhgef12 and active Rho relative to the time period of cell protrusion. Black arrows indicate the time point of maximal sensor or GEF enrichment. The measurements shown in this panel are identical to measurements shown in Figure 4G and Figure 5D. **F**: Schematic representation of the optogenetic strategy to investigate Lbc-type GEF recruitment by active Rac1. **G**: Measurement of Lbc-type GEF and control construct recruitment during acute Rac activation starting at t=0s, n=3 independent experiments with >35 cells per condition. Error bars represent standard error of the mean. Scale bars: 10µm

We next used our new analysis approach to quantify the enrichment dynamics of Arhgef11 and Arhgef12 at the cell edge relative to cell protrusion and retraction events (Figure 5D). This analysis showed that both GEFs were maximally enriched shortly after cell protrusion. By combining measurements of Rac and Rho activity as well as GEF plasma membrane recruitment, we were able to derive a temporal sequence of events (Figure 5E), in which enrichment of active Rac is maximal during cell protrusion with minimal delay (∼0s), followed by the Lbc-type GEFs Arhgef11 (∼160s) and Arhgef12 (∼210 s) and lastly active Rho (>470s).

Similar observations were made using temporal cross-correlation functions (Supplementary Figure 2A). This places maximal Arhgef11 and Arhgef12 enrichment between maximal Rac and Rho activation, which further supports a role for these Lbc-type GEFs in mediating Rac/Rho GTPase activity crosstalk in the coordination of cell protrusion-retraction cycles. To directly investigate the causal relationship between these molecules, we again used the light-controlled PA-Rac1 construct (Wu et al., 2009) and combined optogenetic, acute Rac1 activation with measurements of Arhgef11/12 plasma membrane recruitment. With this approach, we were able to directly show that both Arhgef11 and Arhgef12 are rapidly recruited to the plasma membrane after acute optogenetic Rac1 activation (Figure 5F). Taken together, these results suggest that these GEFs are able to link cell protrusion and retraction by mediating activity crosstalk between Rac and Rho.

### Role of Arhgef11/Arhgef12 in cell protrusion and retraction dynamics

To test, if these GEFs indeed play a role in cell protrusion-retraction cycles and associated cellular processes, we investigated how altering their expression level might affect these dynamic processes. Conceptually, if Arhgef11/12 indeed can mediate Rac/Rho crosstalk, increasing their expression level should increase this crosstalk and thereby shorten the time period that links cell protrusion and retraction. Indeed, ectopic expression of Arhgef11 or Arhgef12 significantly decreased the duration of protrusion-retraction cycles (Figure 6A-B).

**Figure 6:**
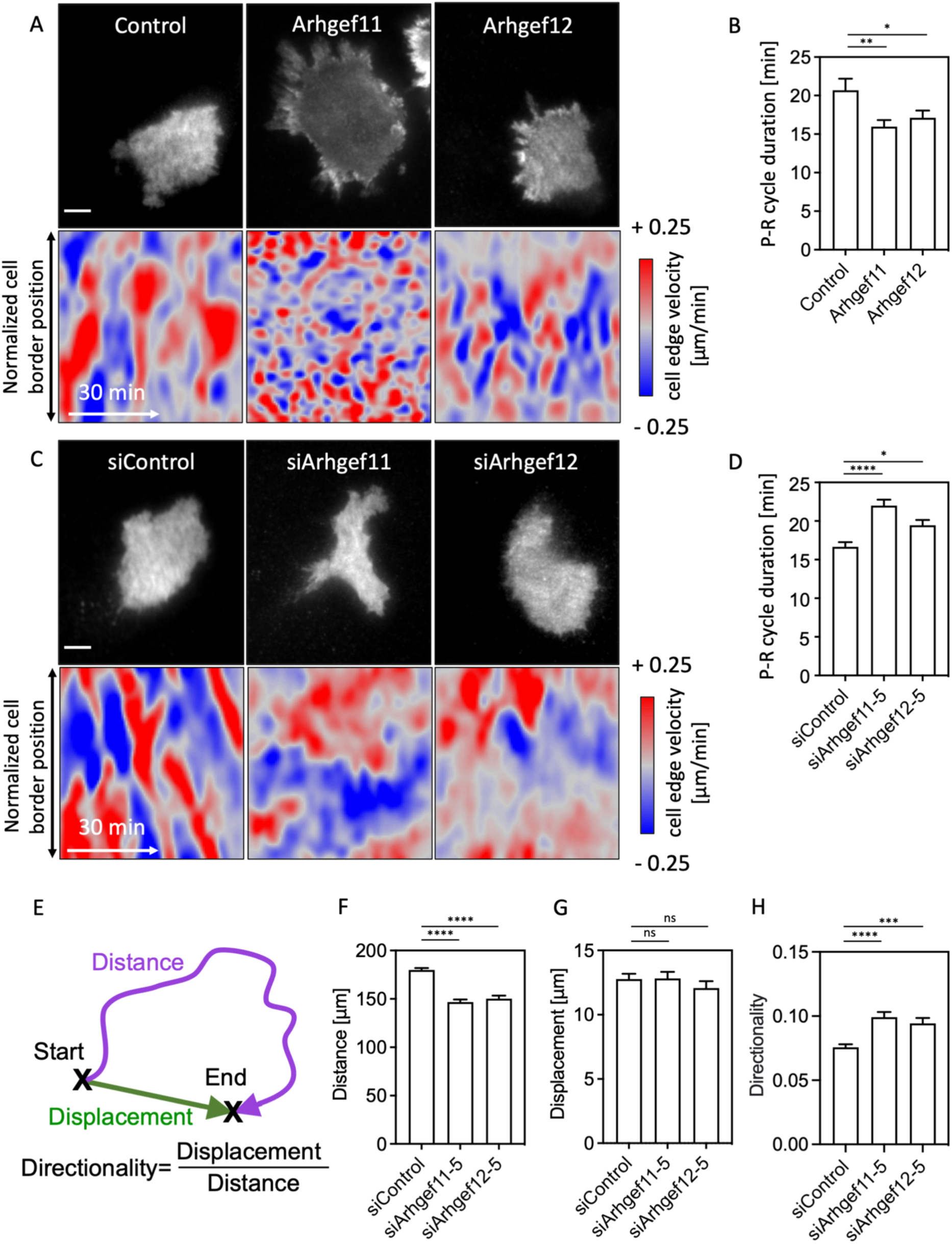
Arhgef11 and Arhgef12 mediate Rac-dependent Rho activation and the spatio-temporal coordination of local cell protrusion-retraction cycles. **A-D**: Quantification of protrusion and retraction dynamics in A431 cells with increased (**A-B**) or decreased (**C-D**) Arhgef11 and Arhgef12 expression levels. (**A,C**): TIRF images (top panels) and cell edge velocity maps (bottom panels) of representative cells that express CMV-mCherry-Arhgef11/Arhgef12 and delCMV-mCitrine (**A**), or Arhgef11/Ahrgef12 targeting siRNA and delCMV-mCherry (**C**). (**B,D**): Quantification of protrusion-retraction (P-R) cycle duration based on cell edge velocity measurements corresponding to panels **A** and **C**, respectively (A-B: n=3 independent experiments with >26 cells per condition, C-D: n=3 independent experiments with >105 cells per condition). **E**: Schematic representation of distance (magenta) and displacement (green) for typical spontaneous exploratory cell migration. The distance corresponds to the length of the cell migration trajectory which leads to the indicated displacement between the start and end locations. The directionality is defined as the ratio between these length measurements. **F-H**: Quantification of distance (F), displacement (G) and directionality (H) of A431 cell trajectories over a 4h time period in control and Arhgef11/Arhgef12 depleted cells (n=3 independent experiments with >491 cells per condition). (*: P<0.05; **: P<0.01; ***: P<0.001; ****: P<0.0001; One-way ANOVA). Images were recorded at a frame rate of 1.5/min (A-D) or 1/min (F-H). Error bars represent standard error of the mean. Scale bars: 10µm.

Conversely, decreasing the expression level of these GEFs would be expected to weaken the link between protrusion and retraction and thus slow down protrusion-retraction cycles. To test this, we used RNA interference to reduce the expression of the endogenous GEFs. As shown by western blot analysis, we were able to efficiently knock down Arhgef11 protein (up to 83±15%) via this strategy and indeed find that this knockdown slowed down protrusion-retraction cycles (Figure 6C-D and Supplementary Figure 2B-C). Knockdown of Arhgef12 was less efficient (up to 75±11%), but the phenotypic effects were similar, although slightly weaker. Taken together, these findings suggest that Arhgef11/12 mediate Rac/Rho activity crosstalk, and that this crosstalk plays a role in the dynamic interplay between cell protrusion and cell retraction dynamics.

### Role of Arhgef11/Arhgef12 in cell migration

Dynamic cell protrusion-retraction cycles are associated with the exploratory cell migration mode typically observed in A431 cells. We therefore investigated, if the frequent cell shape changes that are mediated by Arhgef11/12 contribute to efficient cell migration. Indeed, tracking of individual cells showed that loss of these GEFs significantly decreased migration distance (Figure 6E-F and Supplementary Figure 2D) and therefore reduced their ability to explore their environment. Interestingly, the total displacement was less affected by Arhgef11/Arhgef12 knockdown (Figure 6G) and even increased with one of the Arhgef11 siRNAs (Supplementary Figure 2E), suggesting that the reduced exploratory migration was associated with an increase in migration directionality. Indeed, the ratio of displacement to distance, a typical measure for directionality of migration trajectories, was significantly increased after Arhgef11 or Arhgef12 knockdown (Figure 6E,H and Supplementary Figure 2F).

## Discussion

In this study we investigated the signal crosstalk between the major Rho GTPases Rac, Rho and Cdc42. These investigations surprisingly revealed that acute activation of the cell protrusion regulator Rac1 leads to a strong activity response of the cell contraction regulator Rho. To characterize the function of the Rac/Rho crosstalk in cells, we focused on highly dynamic cell protrusion-retraction cycles that are typically observed during spontaneous, mesenchymal cell migration. Our detailed analysis of Rac and Rho during these cycles revealed that their activities were tightly coupled with phases of cell protrusion and retraction, and the Rac/Rho crosstalk that we measured in our study might play a role in this process. Conceptually, coupling of Rac and Rho activity could be mediated by several distinct mechanisms. Our direct investigation shows that Rac activates Rho, and that conversely, Rho inhibits Rac. Such a link would represent a negative feedback mechanism, which is distinct from the often-cited idea that Rac and Rho mutually inhibit each other (Bolado-Carrancio et al., 2020; Guilluy et al., 2011).

The idea of mutual inhibition between Rac and Rho appears quite attractive, as it can be used to explain cell polarization of migrating cells, in which a Rac-dependent protrusive front region and a Rho-dependent contractile back region mutually exclude each other. Indeed, a mechanism based on mutual inhibition alone is expected to stably segregate protrusive front and contractile back signals as it is observed in persistent cell migration of neutrophils (Wong et al., 2006). However, additional mechanisms would be required to enable dynamic cycling of protrusive and contractile signals as it is observed during exploratory migration of mesenchymal cells (Nalbant & Dehmelt, 2018). The oscillatory or excitable system dynamics observed for protrusive front and contractile back signals could play a role in the formation of such cycles of protrusion and retraction. However, to coordinate these cycles, front and back signals have to be coupled with each other. The asymmetric hierarchy, which is implied by our crosstalk analysis, i.e. activation of Rho by Rac and inhibition of Rac by Rho, is expected to enable the temporal cycles of protrusions followed by retractions, which are typically observed in such cells. A signal network that is based on mutual inhibition between Rac and Rho instead would lock a local cell area either in a protrusive or retractile state and would require an additional signal to dynamically switch local cell shape changes.

While the concept of mutual inhibition between Rac and Rho is frequently cited, several earlier studies suggested a potential activating role of Rac on Rho (Bustos et al., 2008; Guilluy et al., 2011; Nimnual et al., 2003; Rosenfeldt et al., 2006; Sander et al., 1999). Most prominently, initial studies of cellular Rho GTPase function showed that injection of an active Rac mutant induced formation of stress fibers that were dependent on Rho activity (Ridley et al., 1992). To investigate this mechanism further, we focused on the recent finding that members of the Lbc family of Rho GEFs might mediate Rac-dependent Rho activation (Dada et al., 2017). By combining acute optogenetic Rac activation with TIRF microscopy, we indeed identified both Arhgef11 and Arhgef12 as Rac effectors, and both molecules are localized to the peripheral edge of cells. This suggests that these molecules might be able to coordinate dynamic cell shape changes associated with Rac and Rho activity. Increasing or decreasing the expression level of either Arhgef11 or Arhgef12 revealed that these molecules indeed play a role in the coupling between dynamic cell protrusions and retractions.

Together with previous work, we propose a mechanism, in which Arhgef11/12 couple the signal modules that control cell protrusion and cell retraction (Figure 7). We showed that active Rac can recruit Arhgef11/12 (Figure 5F), which are well-known to activate Rho (Fukuhara et al., 1999, 2000). Increased Rho activity can be further amplified via positive feedback (Graessl et al., 2017), and can inhibit Rac activity (Figure 2E-F). Rho activity itself can be inhibited via negative feedback (Graessl et al., 2017) to enable another protrusion-retraction cycle.

**Figure 7:**
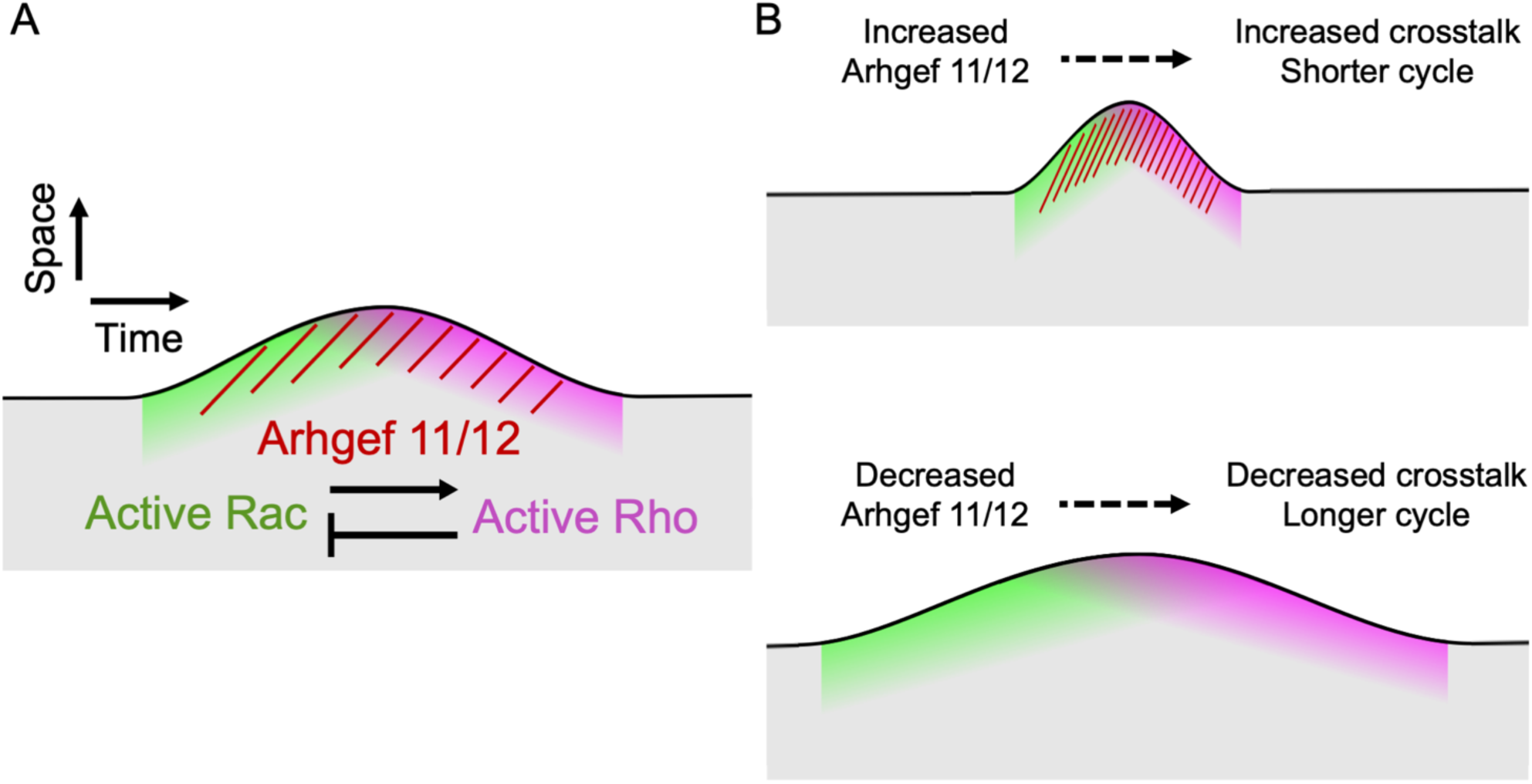
Proposed mechanism for the generation of protrusion retraction cycles. **A**: Schematic for spatio- temporal events that couple cell protrusion and retraction and the signal molecules that mediate this coupling. **B**: Effect of increasing or decreasing Arhgef11/12 levels on local Rac/Rho crosstalk and cell morphodynamics.

Finally, our study revealed how Arhgef11/12 are linked to exploratory cell migration. By coupling Rac/Rho activity, Arhgef11/12 are expected to mediate cell retraction after protrusion and thus facilitate spontaneous changes in migration direction. Indeed, we find that knockdown of Arhgef11/12 increased directionality during cell migration (Figure 6G). Thus, our study reveals a mechanism, how crosstalk between the major cell morphogenesis regulators Rac and Rho links local cell protrusion and retraction dynamics to enable effective exploratory cell migration.

## Experimental Procedures

### Cell culture and reagents

Neuro-2a cells (DSMZ, Braunschweig) were cultured in MEM Eagle (10% FBS, 100 U/ml Penicillin + 100 µg/ml Streptomycin, 2 mM L-Glutamine, 1 mM Sodium Pyruvate, PAN Biotech). A431 cells (ATCC) were cultured in DMEM medium (10%FBS and 2 mM L-Glutamine, PAN Biotech). HeLa, NIH3T3, and U2OS cells were cultured in DMEM medium (10%FBS and 50U/ml Penicillin + 50 µg/ml Streptomycin, 2 mM L-Glutamine, PAN Biotech). All cells were maintained using standard culture techniques at 37°C and 5% CO_2_. For live-cell imaging, Neuro-2a, HeLa, and NIH3T3 cells were plated onto LabTek glass surface slide (Thermo Fischer Scientific) or glass bottom dishes (MatTek) before transfection of plasmid DNA. Pre-treatments of glass surfaces were as follows: uncoated for Neuro-2a and Hela cells, poly-L-lysine for NIH3T3 cells and 10 ug/ml Collagen-I for U2OS cells. A431 cells were plated on LabTek glass surfaces that were coated with 10 ug/ml Fibronectin for 45 min. The SLF’-TMP dimerizer and TMP competitor were synthesized as described previously (Liu et al., 2014). To quantify GEF/sensor enrichment at the cell periphery, cells were plated on 35 mm culture dishes, transfected with plasmid DNA, and then replated on fibronectin-coated MatTek glass bottom dishes (10 ug/ml fibronectin for 16 hours at 4°C).

### Plasmid Constructs and siRNA

EGFP-2x-FKBP’-Rac1Q61LΔCAAX and TagBFP-2xeDHFR-CAAX for targeting active Rac1 to the plasma membrane via chemically-induced dimerization were described previously (Liu et al., 2014). Here, optimized constructs for targeting of Rac1, Cdc42 and RhoA were prepared. First, EGFP in EGFP-2x-FKBP’-Rac1Q61LΔCAAX was replaced by mTurquoise2 to facilitate simultaneous measurement of the four fluorophores TagBFP, mTurquoise2, mCitrine and mCherry. In addition, a nuclear export sequence was introduced between mTurqouise2 and FKBP’ to prevent nuclear accumulation, which results in delayed plasma membrane recruitment. In brief, mTurquoise2-2xFKBP’-Rac1Q61LΔCAAX was generated by ligating the larger fragment from EGFP-2xFKBP’-Rac1Q61LΔCAAX (Liu et al., 2014) with the smaller fragment from pmTurquoise2-N1 (Addgene Plasmid #60561), after digestion with BsrGI and NheI. In a second step, this resulting plasmid and a PCR fragment amplified from pmTurquoise2-NES (Goedhart et al., 2012) (Addgene Plasmid# 36206) with primers 5’-TCAGTTGCTAGCCTCAAGCTTCGAATTCTG-3’ and 5’-AGAGTCAGCTCGAGATATCTTGTACGAGTCCAG-3’, were digested using NheI/XhoI. The larger fragment of the plasmid was ligated to the PCR product to yield the final perturbation construct mTurquoise2-NES-2xFKBP’-Rac1Q61LΔCAAX. The analogous Cdc42 and RhoA perturbation constructs were generated as derivatives from this Rac1 construct by ligation to PCR fragments amplified from pcDNA3-EGFP-Cdc42Q61L or pcDNA3-EGFP-RhoAQ63L (kind gifts from Gary Bokoch, The Scripps Research Institute) using 5’-CTGTACTCTAGATCCATGCAGACAATTAAGTG-3’/5’-TCGAGTCAATTGAGTTAGGACCTGCGGCTCTTC-3’ and 5’- GGAATTCTAGATCCATGGCTGCCATCCGGAAG-3’/5’-CGAGTCAATTGAGTTAGGAACCAGATTTTTTC-3’, respectively, after digestion of both fragments using MfeI/XbaI. The plasmid coding for mCherry fused to β-actin, driven by the Ubiquitin-C promotor (mCherry-actin-Ub) was generated in two steps: 1. mCherry-C1-Ub was generated from pUB-GFP (Addgene plasmid #11155) by inserting mCherry amplified from mCherry-C1 (Clontech) using 5’-CGGCATTAATGATCTGGCCTCCGCGCCGGGT-3’ and 5’-GGCCGCTAGCCGACCTGCAGCCCAAGCTTCGTC-3’, after digestion of both fragments with AseI/NheI, 2. mCherry-actin-Ub was generated from mCherry-C1-Ub by inserting β-actin amplified from mRFP1-actin (Dehmelt et al., 2006) using 5’- ATATGAATTCCGCCCCGCGAGCACAGA-3’ and 5’-ATATGGATCCTCAGTGTACAGGTAAGCCCTGGC-3’, after digestion of both fragments with EcoRI/BamHI. The Rac, Cdc42 and Rho activity sensor constructs driven by the low-expressing delCMV promotor (Watanabe & Mitchison, 2002), delCMV-mCherry-p67^phox^-GBD, delCMV-WASP-GBD and delCMV-mCherry-Rhotekin-GBD were described previously (Graessl et al., 2017). The control construct delCMV-mCitrine was generated by ligating the larger fragment from pmCitrine-N1 (Addgene Plasmid #54594) with the smaller fragment from the delCMV-mCitrine-RBD biosensor (Graessl et al., 2017), after digestion with AseI and BsrGI. delCMV-mCherry was generated by ligating the larger fragment from pmCherry-N1 (Clontech) with the smaller fragment from delCMV-mCherry-actin (Schulze et al., 2014), after digestion with AseI and BsrGI. mCerulean-PA-Rac1Q61L was a kind gift from Klaus Hahn (University of North Carolina). mCherry-NMHCIIA (Dulyaninova et al., 2007) was obtained from Addgene (Plasmid #35687). The majority of transfections were performed using Lipofectamine™2000 (Thermo Fisher Scientific). Transfections of A431 cells were performed using Lipofectamine™3000 (Thermo Fisher Scientific). For initial experiments in Neuro-2a cells (Figure 1), XtremeGene 9 (Roche Diagnostics) was used.

The improved Rac1 activity sensor (delCMV-mCherry-3X-p67^phox^-GBD) was generated from the previously established delCMV-mCherry-p67^phox^ -GBD construct (Graessl et al., 2017). The sensor was generated in two steps via PCR based Gibson assembly. For the Gibson reaction, a mix was prepared consisting of 5 % PEG-8000 (Promega), 100 mM Tris-HCl (pH 7.5), 10 mM MgCl2, 10 mM DTT, 0.2 mM dATP, 0.2 mM dTTP, 0.2 mM dCTP, 0.2 mM dGTP, 1 mM NAD (New England Biolabs), 2.0 U T5 exonuclease (New England Biolabs), 12.5 U Phusion DNA polymerase (New England Biolabs), 2000U Taq DNA ligase (New England Biolabs). First, the p67^phox^-GBD insert was amplified using 5’- GGACTCAGATCTCGAGCTCACATGTCCCTGGTGGAGGCCA-3’/5’- ACCAGGGACATGGAATTCGATCCACTTCCAGAACCCGTCGCCTTGCCTAGGTAATC-3’ and inserted into the parent plasmid after linearization using HindIII to generate delCMV-mCherry-2x-p67^phox^- GBD. In the second step, one p67^phox^-GBD repeat was amplified from delCMV-mCherry-2x-p67^phox^-GBD using 5’-ACGGGTTCTGGAAGTGGATCGGTTCTCATGTCCCTGGTGGAGGC-3’/5’-GGCCTCCACCAGGGACATGGAATTCGATCCACTTCCAGAACCCGTCGCCTTGCCTAGGTAATC-3’ and inserted into the EcoRI-cut delCMV-mCherry-2x-p67^phox^-GBD plasmid to generate delCMV-mCherry-3x-p67^phox^-GBD. The Rho activity sensor (delCMV-mCherry-2X-RhotekinGBD) was generated using the previously established delCMV-mCherry-RhotekinGBD construct (Graessl et al., 2017). The sensor was generated using an analogous PCR based Gibson assembly as described above. First 1x-RhotekinGBD was amplified using 5’-TACAAGTCCGGACTCAGATCTCGAGAAGCTTCGAATTCCCTGG-3’/5’-AGGGAATTCGAAGCTTGAGCGAGTCCGGAGCCTGTCTTCTCCAGCAC-3’. The amplified fragment was then inserted into the at XhoI-cut parent plasmid.

CMV promoter-driven Lbc GEF encoding plasmids (CMV-mCherry-Arhgef1, CMV-mCherry-Arhgef11, CMV-mCherry-Arhgef12, CMV-mCherry-AKAP13, CMV-mCherry-Arhgef18, and CMV-mCherry-Arhgef28) were gifts from Oliver Rocks (Müller et al., 2020). CMV-mCherry-Arhgef2 was generated using PCR based Gibson assembly by replacing Arhgef1 in the CMV-mCherry-Arhgef1 plasmid with Arhgef2. First, Arhgef2 (also known as GEF-H1) was amplified from the previously established EGFP-GEF-H1 (Krendel et al., 2002) plasmid using 5’- AGTCCGGACTCAGATCTCGAGGGCGCGCCATGTCTCGGATCGAATCCC-3’/5’-GATCCGGTGGATCCTTAGTTAATTAAGCTCTCGGAGGCTACAGC-3’. The amplified fragment was then inserted into CMV-mCherry-Arhgef1, after removing the Arhgef1 insert flanked by AscI and PacI sites.

A delCMV promoter-driven Arhgef1 encoding plasmid (delCMV-mCherry-Arhgef1) was generated using Gibson assembly by amplifying the protein coding sequence from CMV-mCherry-Arhgef1 plasmid using 5’-CTCCACCGGCGGCATGGACGAGCTGTACAAGTCCGGACTCAGATCTC 3’/5’TCTAGAGTCGCGGCCGCTTTACTTTTAGTTAATTAAAGTGCAGCCAG 3’. The amplified fragment was then inserted into a BsrGI cut delCMV-mCherry plasmid. delCMV-mCherry-Arhgef1 was then used to generate delCMV-mCherry-Arhgef11 by replacing Arhgef1 with Arhgef11 from CMV-mCherry-Arhgef11 at AscI/PacI sites. delCMV-mCherry-Arhgef12 was generated using Gibson assembly by amplifying Arhgef12 from CMV-mCherry-Arhgef12 using 5’-GCTGTACAAGTCCGGACTCAGATCTCGAGGGCGCGCCAGTGG-CACACAGTCTACTATC-3’/5’-CCGCTTTACTTTTAGTTAATTAAACTTTTATCTGAGTGCTTGTC-3’. The amplified fragment was then inserted into delCMV-mCherry-Arhgef1, after removing the Arhgef1 insert flanked by BglII and PacI.

For knockdown experiments, ON-Target plus siRNAs (Dharmacon^TM^) were used (siControl: **#2** 5’-UGGUUUACAUGUUGUGUGA-3’, siArhgef11: **#5** 5’-GCAAGUGGCUGCACAGUUC-3’, **#7** 5’-UCUAUGAGCUGGUUGCAUU-3’, siArhgef12: **#5** 5’-GAUCAAAUCUCGUCAGAAA-3’, **#6** 5’-GAAAUGAGACCUCUGUUAU-3’). Briefly, cells were transfected with 30nM of the siRNAs using Lipofectamine^TM^ RNAiMAX (Invitrogen). Cells were incubated with the siRNAs for 48 hours before splitting and seeding for experiments. Experimental analysis of knockdown phenotypes was performed 96 hours post siRNA treatment and quantification of knockdown efficiency was performed at that same timepoint via Western blot analysis.

### Western blot analysis

Cells were washed one time with ice cold PBS, then lysed with ice cold 1x cell lysis buffer (9803, CST) for 5 min on ice. The cell lysate was then centrifuged at 13000 rpm for 10 min at 4°C to remove insoluble material. A Bradford assay was used to measure the protein concentration in the supernatant. 5x Laemmli sample buffer was used to prepare protein samples, which were boiled at 95°C for 5 min before being separated using SDS-PAGE (4561086, Biorad).

Wet blot transfer was used to transfer proteins to a PVDF membrane (MERCK). Intercept blocking buffer (927-60001, LI-COR) was used to block the membrane for 60 min at room temperature.

Blots were incubated for 24 h with primary antibodies at 4°C while shaking (Arhgef 12 antibody GTX87286 at 1:1000; Arhgef11 antibody sc-166740 at 1:50; GAPDH antibody CST-2118 as loading control at 1:1000). All antibodies were diluted in intercept blocking buffer. The membranes were then washed with TBS-T buffer and stained with secondary antibodies (926-68070, 926-32211, IRDye®, Licor at 1:10,000) for 60 min at room temperature. After final washing steps, the blots were measured using the Odyssey® CLx imaging system (LI-COR).

### Microscopy

TIRF microscopy was performed on an Olympus IX-81 microscope, equipped with a TIRF-MITICO motorized TIRF illumination combiner, an Apo TIRF 60×1.45 NA oil immersion objective and a ZDC autofocus device. For the majority of experiments that employed spectral emission ranges in blue (TagBFP), cyan (mTurquoise2/mCerulean), yellow (mCitrine) and red (mCherry), a dichroic mirror (ZT405-440/514/561) was used in combination with an emission filter set (HC 435/40, HC 472/30, HC 542/27 and HC 629/53), the 514 nm OBIS diode laser (150 mW) (Coherent, Inc., Santa Clara, USA) or the 514 nm line of a 400 mW Argon ion laser (model # 543-A-A03, Melles Griot, Bensheim, Germany), and the Cell R diode lasers (Olympus) with wavelength 405nm (50 mW), 445nm (50 mW) and 561nm (100 mW), as well as wide-field illumination via the MT20 light source (Olympus) or the Spectra X light engine (Lumencor). For detection, this was combined with an EMCCD camera (C9100-13; Hamamatsu, Herrsching am Ammersee, Germany) at medium gain without binning. The microscope was equipped with a temperature-controlled incubation chamber. Time-lapse live-cell microscopy experiments were carried out with indicated frame rates at 37°C in CO_2_-independent HEPES-stabilized imaging medium (Pan Biotech) supplemented with 10% FBS.

Tracking of single cell migration was performed using an Olympus IX81 microscope with a UPlanSApo 10x objective. Wide-field images were acquired using a 651 nm LED lamp with Spectra X light engine (Lumencor). The microscope was equipped with a temperature-controlled incubation chamber. Time-lapse live-cell microscopy experiments were carried out at 37°C in CO_2_-independent HEPES-stabilized imaging medium (Pan Biotech) supplemented with 10% FBS.

### Perturbation via chemically-induced dimerization

Chemically-induced dimerization was performed essentially as described before (Liu et al., 2014). Briefly, synthesis and purification of the dimerizer SLF’-TMP and of the competitor TMP followed established protocols (Liu et al., 2014). Neuro-2a cells were transfected with TagBFP-2xeDHFR-CAAX and mTurquoise2-NES-2xFKBP’ fused to the Q61LΔCAAX mutant of Rac1, Cdc42 or RhoA. To investigate morphological changes, cells were co-transfected with mCherry-actin-Ub, to investigate Rho GTPase crosstalk, cells were co-transfected with the Rac, Cdc42 or Rho sensor constructs delCMV-mCherry-p67^phox^-GBD, delCMV-WASP-GBD or delCMV-mCherry-Rhotekin-GBD, respectively. In all experiments, the delCMV-mCitrine control sensor was co-expressed. Chemically-induced dimerization was initiated by addition of 10 μl SLF’-TMP dimerizer and stopped by addition of 10 μl TMP competitor.

### Optogenetic perturbation

Cells were transfected with photo-activatable Rac1 (mCerulean-PA-Rac1) and light-based activation of mCerulean-PA-Rac1 was performed by TIRF illumination using the 445nm Cell R diode laser. To prevent saturated PA-Rac1 activation, 1000x-10000x neutral-density filters were added into the 445nm TIRF illumination light path. The built-in neutral-density filter wheel was used to fine-tune light intensity and was typically set to ∼30%. Within photoactivation time intervals, 445nm TIRF illumination was constantly on, except for the exposure times during image acquisition. To minimize background activation of mCerulean-PA-Rac1, illumination intensity, duration and frequency were kept as low as possible and fluorescence measurements were always performed using the 514nm and 561nm excitation lines, including the excitation of EGFP fluorophores. Detection of TagBFP, mTurquoise2 or mCerulean was always performed after the experiment.

### Analysis of subcellular morphodynamics

To investigate the local enrichment of Lbc type GEFs in cell protrusions and retractions, A431 cells were transfected with plasmids coding for mCherry fused Lbc GEFs driven by the CMV promotor. To improve the signal-to background ratio, the automated quantification of Arhgef11 and Arhgef12 signals at the cell periphery was performed by transfecting mCherry-GEF constructs driven by the delCMV promoter. To study the local enrichment of the GTPase activity in cell protrusions and retractions, cells were transfected with the enhanced Rho or Rac sensor. To study the effect of overexpression of Arhgef11 and Arhgef12 on cell morphodynamics, cells were transfected with plasmids coding for mCherry fused GEFs driven under CMV promoter. In all experiments described above, delCMV-mCitrine was co-transfected to identify the cell edge independent of the signals of interest. To study the effect of siRNA mediated knockdown on cell morphodynamics, delCMV-mCherry was used to identify the cell borders.

### Analysis of cell migration

The effect of siRNA knockdown on cell migration was investigated by staining nuclei in living A431 cells using the SPY650-DNA dye (Spirochrome). 96 hours post siRNA treatment, cells were seeded on 10μg/ml fibronectin coated LabTek dishes at a density of ∼143 cells/mm^2^. 6 hours post seeding, cells were incubated with SPY-650 (1:1000) in imaging medium for 1.5 h. Cells were then washed once with fresh imaging media and rested for 1 h before onset of imaging. Fluorescence measurements were performed using the 651 nm LED lamp of the SpectraX light engine. Images were collected with a frame rate of 1/min.

### Data analysis

All image analysis was performed using ImageJ (http://imagej.nih.gov/ij/) and all figures were prepared using Photoshop CS4. Kymograph analysis was performed using the ImageJ built-in multi kymograph plugin. Cell tracking was performed using the Trackmate plugin. Statistical analysis, curve fitting (mono-exponential decay) and generation of plots was performed using Prism (GraphPad).

#### Analysis of Rho GTPase activity crosstalk via chemically induced dimerization

The fluorescence intensity of Rho activity sensor *I_Rho_* was measured via TIRF microscopy in central cell regions that were completely adherent during the entire observation period. During the time course of perturbation via chemically-induced dimerization, cell shrinkage or expansion occurred that lead to Rho activity independent changes in fluorescence intensity, presumably due to changes in cell volume and associated changes in protein concentration. To correct for these intensity changes, we co-expressed a control sensor (delCMV-mCitrine) and measured the fluorescence intensity of the control sensor *I_Control_* in identical cell regions as the intensity of the Rho activity sensor. These raw intensity measurements were normalized by subtracting the background signals outside cell areas *I_Rho,BG_* and *I_Control,BG_* and dividing by the initial, background-corrected intensity values *I_Rho,0_*‒*I_Rho,0,BG_* and *I_Control,0_*‒*I_Control,0,BG_* before the perturbation. The corrected activity sensor measurements *A_Rho,corr_* were obtained by subtracting the normalized control sensor measurements from the normalized Rho GTPase activity sensor measurements:

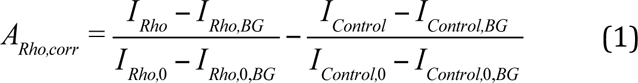

#### Analysis of Rac/Rho GTPase activity crosstalk via optogenetic perturbations

Optogenetic perturbation of Rac1 induced only minor changes in the control sensor signal during the short time periods of the crosstalk measurements (see Figure 3). Therefore, subtraction of control sensor measurements was not necessary in these experiments and activity sensor measurements *A_Rho_* were calculated via the following simplified equation:

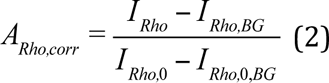

In these experiments, delCMV-mCitrine or delCMV-mCherry were used as control sensors, which were either co-expressed with the Rho sensor (N2A, NIH3T3, Hela or U2OS cells), or expressed in a separate cell population (A431 cells).

#### Analysis of cell morphodynamics and local fluorescence signals at the cell edge

Analysis of local cell edge velocity and associated local fluorescence signals were performed using a modified version of the ADAPT plugin (Barry et al., 2015) in combination with custom ImageJ analysis scripts. In brief, the published ADAPT plugin performs analysis of local signal intensities via a region that extends from the cell edge both inwards, in the direction of the cell center and outwards, away from the cell. Thereby, signals that are localized near the cell edge are decreased by averaging with the local background intensity and therefore do not permit quantitative estimations of relative, local signal enrichment. To enable such estimations, the ADAPT code was modified to generate a region that only extends from the cell edge inwards. The signal and cell velocity maps and signal/velocity crosscorrelation functions were directly extracted from the standard output of the modified ADAPT script. To calculate the local signal enrichment, spatio-temporal regions in the cell velocity maps that correspond to cell protrusion (>0.075μm/min) or retraction (< −0.075μm/min) were selected via the threshold function and transferred to the fluorescence signal maps. The average intensity within these regions was measured and the enrichment was calculated as the ratio of this intensity divided by the average intensity in the whole cell (including the cell edge and central cell regions) over the full time period. To generate the local signal enrichment functions, the spatio-temporal regions described above were shifted along the temporal x-axis of the fluorescent signal map, to extract the time-shifted local signal enrichment. To calculate the duration of protrusion-retraction cycles, the cell periphery was divided into 100 positions. This was achieved by rescaling the cell velocity maps in the position axis to a size of 100 pixels. In each horizontal line of the rescaled cell velocity maps, time periods were identified as protrusions or retractions. Protrusion-retraction cycle duration was defined as the time period starting from the onset of a protrusion and the following onset of a retraction. For experiments to investigate the effect of increasing Arhgef11/12 levels via ectopic expression or decreasing Arhgef11/12 levels via RNAi, the threshold to identify protrusions or retractions was the same as for the signal enrichment analysis (>0.075μm/min for protrusions and <-0.075μm/min for retractions).

#### Analysis of cell migration velocity and directionality

Quantification of velocity and directionality was performed using the TrackMate plugin (Tinevez et al., 2017) in ImageJ. SPY-650 stained nuclei were selected and segmented using an intensity threshold and size filter. Cells that leave the field of view during tracking were not included in the analysis. Tracks generated by TrackMate were used to calculate velocity and directionality measurements using the chemotaxis plugin (Ibidi GmbH, Martinsried, Germany) in ImageJ.

### Author contributions

L.D., P.N., S.N., A.C, and T-T.D. designed the research. S.N. performed, analyzed and optimized the majority of experiments to investigate protrusion-retraction dynamics in A431 cells and the role of GEFs in mediating Rac/Rho crosstalk. T-T.D. performed, analyzed and optimized the majority of combined Rho activity perturbation-response and effector activation measurements. A.C. developed, characterized, and optimized the activity perturbation and the activity measurements and performed initial Rac1-Rho crosstalk analyses. S.N. contributed to characterization of activity perturbation method development. S.N. developed improved Rac and Rho activity sensors based on tandem GBDs. J.K. performed initial combined perturbation-response analysis and contributed to combined perturbation-response analysis. A.S. contributed to investigation of GEFs in mediating Rac/Rho crosstalk. D.S. contributed to activity sensor development. Y.W.W. and X.X. contributed to activity perturbation development and synthesized the chemical dimerizer. L.D. and P.N. supervised experiments. L.D., and P.N. wrote the majority of the manuscript. All authors contributed to discussions and manuscript preparation.

## Acknowledgements

We thank Sven Müller (MPI Dortmund) for expert microscopy support and Philippe Bastiaens (MPI Dortmund) for departmental support and helpful discussions. This work was supported by MERCUR grant No. Pr-2012-0022 to P.N. and L.D., FORSYS partner initiative (BMBF grant No. 0315258) to L.D., the Deutsche Forschungsgesellschaft DFG project grant 823/3-1 to L.D. and S.N., DFG Heisenberg Programme grant 823/4-1 to L.D., DFG Principal Investigator grant DE 823/10-1 to L.D., a DAAD Ph.D. fellowship to A.S., the European Research Council, ERC (ChemBioAP) to Y.W.W., the Knut and Alice Wallenberg Foundation to Y.W.W., the Göran Gustafsson Foundation for Research in Natural Sciences and Medicine to Y.W.W. and Vetenskapsrådet (Nr. 2018-04585) to Y.W.W.

## Supplementary Figures

**Supplementary Figure 1:**
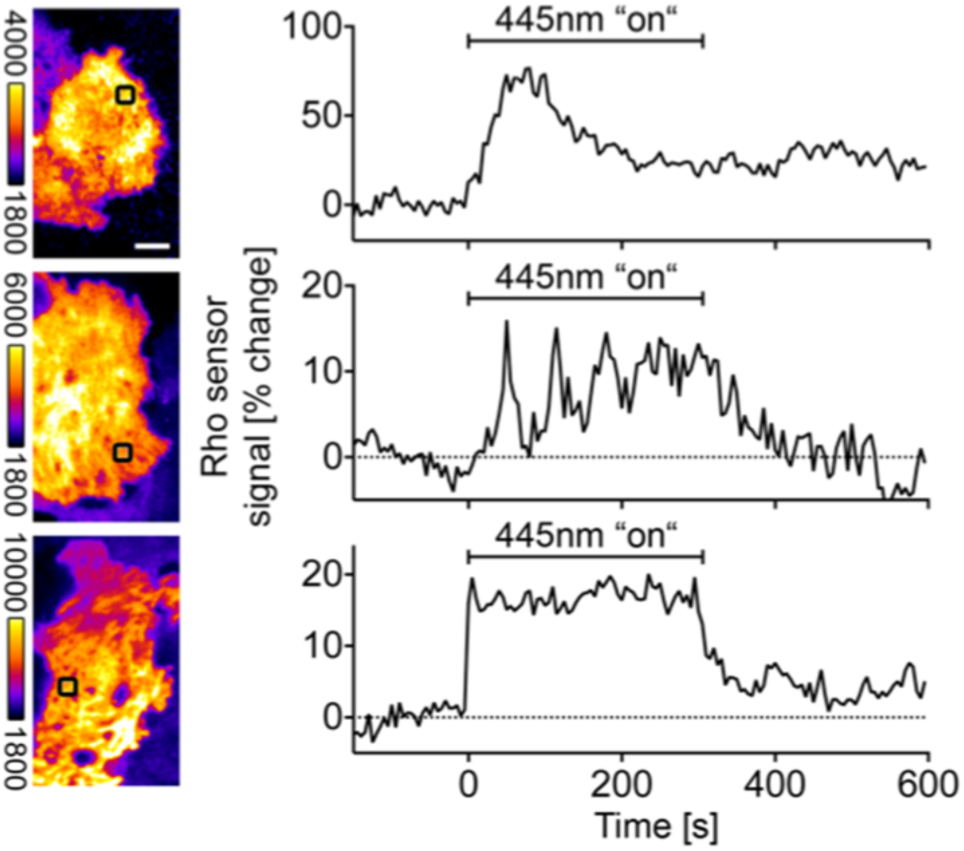
Dynamic Rho activity response to acute Rac activity perturbation. Examples of Rac1-induced Rho activity dynamics in individual U2OS cells. Numbers are indicated as percentage ± standard error of the mean. Left: Representative TIRF images. Right: Local Rho activity sensor measurements corresponding to black boxes in the TIRF images on the left. 56±11% of 44 cells showed a discernible Rho activity response. Top graph: transient single pulse response (46±30% of the 26 responding cells), Middle graph: increased pulse frequency/amplitude (24±12% of the 26 responding cells), Bottom graph: general Rho activity increase (30±20% of the 26 responding cells). All observations and measurements are based on at least 3 independent repetitions. Scale bar: 10 µm.

**Supplementary Figure 2:**
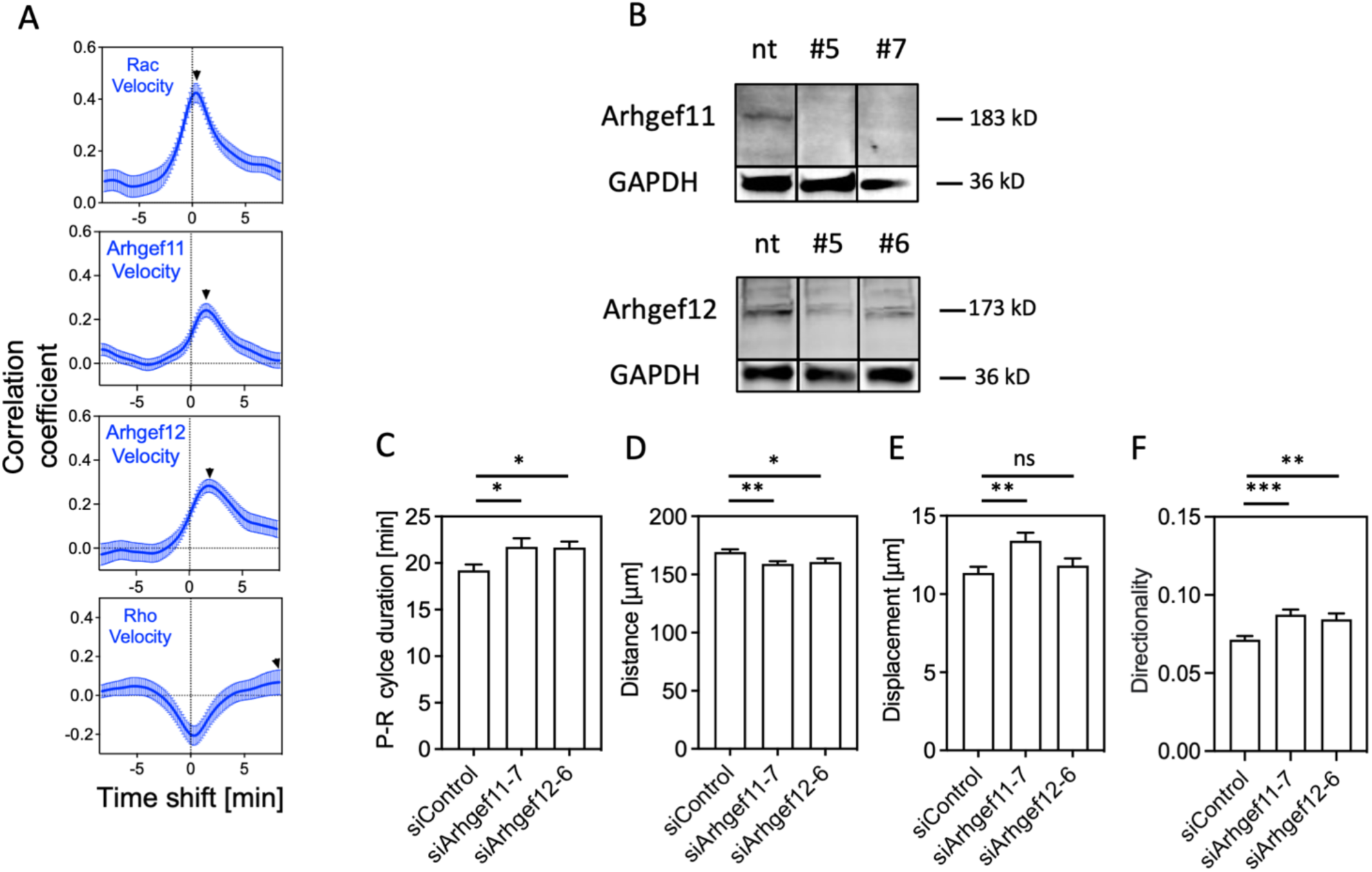
Investigation of the role of Arhgef11 and Arhgef12 in Rac Rho activity crosstalk, cell morphodynamics and cell migration. **A:** Direct comparison of signal-cell edge velocity crosscorrelation functions for active Rac, Arhgef11, Arhgef12 and active Rho, related to Figure 5E. Black arrows indicate the time point of maximal sensor or GEF correlation. The measurements shown in this panel are identical to measurements shown in Figure 4F and Figure 5D. **B**: Quantification of Arhgef11/12 knockdown via Western blot analysis. A representative blot is shown (n=3 independent repetitions). Quantification of knockdown efficiency: 83±15% for Arhgef11-5, 75±11% for Arhgef12-5, 80±10.3% for Arhgef11-7 and 70±16.7.% for Arhgef12-6 (percent ± standard error of the mean). **C**: Quantification of protrusion-retraction (P-R) cycle duration based on cell edge velocity measurements related to Figure 6D using a second set of siRNA oligonucleotides (n=3 independent experiments with >158 cells per condition). **D-F**: Quantification of distance (D), displacement (E) and directionality (F), of A431 cell trajectories over a 4h time period in control and Arhgef11/Arhgef12 depleted cells, related to Figure 6F-H, using a second set of siRNA oligonucleotides (n=3 independent experiments with >484 cells per condition). (*: P<0.05; **: P<0.01; ***: P<0.001; One-way ANOVA). Images were recorded at a frame rate of 1.5/min (C) or 1/min (D-F). Error bars represent standard error of the mean.

## Supplementary Movies

Movie 1: **Morphological changes during acute and reversible perturbation of the small GTPases Rac1, Cdc42 and RhoA via chemically-induced dimerization (related to** Figure 1B). Time-lapse TIRF videos of mCherry-Actin in representative Neuro-2a cells before and during the application of the chemical dimerizer SLF’- TMP and the competitor TMP, which induce reversible plasma membrane targeting of dominant positive mutants of the small GTPases Rac1, Cdc42 and RhoA. Images were collected with a frame rate of 2/min.

Movie 2: **Direct investigation of Rac1-Rho crosstalk in living cells via chemically-induced dimerization (related to** Figure 2B). Time-lapse TIRF videos of the perturbation construct mTurquoise2-Rac1Q61LΔCAAX, the Rho activity sensor mCherry-Rhotekin-GBD and the control sensor mCitrine in a representative Neuro-2a cell before and during the application of the chemical dimerizer SLF’-TMP and the competitor TMP. Images were collected with a frame rate of 3/min.

Movie 3 and 4: **Rac activity dynamics in protrusion-retraction cycles in spontaneously migrating A431 cells (related to** Figure 4A-E). Time-lapse TIRF videos of the improved Rac activity sensor (mCherry-3xp67^phox^GBD; green) in A431 cells. A cytosolic cell filler (mCitrine) was co-expressed to detect the cell attachment area (magenta). Images were collected with a frame rate of 6/min.

Movie 5: **Rho activity dynamics in protrusion-retraction cycles in spontaneously migrating A431 cells (related to** Figure 4E). Time-lapse TIRF videos of the improved Rho activity sensor (mCherry-2X-RhotekinGBD; green) in a A431 cell. A cytosolic cell filler (mCitrine) was co-expressed to detect the cell attachment area (magenta). Images were collected with a frame rate of 6/min.

Movie 6: **Arhgef11 plasma membrane association dynamics in protrusion-retraction cycles in spontaneously migrating A431 cells (related to** Figure 5C). Time-lapse TIRF videos of a fluorescently labeled Arhgef11 construct (mCherry-Arhgef11; green) in a A431 cell. A cytosolic cell filler (mCitrine) was co-expressed to detect the cell attachment area (magenta). Images were collected with a frame rate of 6/min.

Movie 7: **Arhgef12 plasma membrane association dynamics in protrusion-retraction cycles in spontaneously migrating A431 cells (related to** Figure 5C). Time-lapse TIRF videos of a fluorescently labeled Arhgef12 construct (mCherry-Arhgef12; green) in a A431 cell. A cytosolic cell filler (mCitrine) was co-expressed to detect the cell attachment area (magenta). Images were collected with a frame rate of 6/min.

